# NEAT1 involves Alzheimer’s Disease (AD) progression via regulation of glycolysis and P-tau

**DOI:** 10.1101/643718

**Authors:** Yiwan Zhao, Ziqiang Wang, Yunhao Mao, Bing Li, Yuanchang Zhu, Shikuan Zhang, Songmao Wang, Yuyang Jiang, Naihan Xu, Yizhen Xie, Weidong Xie, Yaou Zhang

**Affiliations:** School of Life Sciences, Tsinghua University, Beijing 100084, P.R. China; Key Lab in Healthy Science and Technology, Division of Life Science, Graduate School at Shenzhen, Tsinghua University, Shenzhen 518055, P.R. China; State Key Laboratory of Chemical Oncogenomics, Graduate School at Shenzhen, Tsinghua University, Shenzhen, P.R. China; Key Laboratory of Medical Reprogramming Technology, Shenzhen Second People’s Hospital, First Affiliated Hospital of Shenzhen University, Shenzhen 518035, China.; Open FIESTA Center, Tsinghua University, Shenzhen 518055, P.R. China.; State Key Laboratory of Applied Microbiology Southern China, Guangdong Provincial Key Laboratory of microbial Culture Collection and Application, Guangdong Institute of microbiology, Guangdong, 510070, P.R. China.

**Keywords:** Alzheimer’s Disease, NEAT1, FZD3, H3K27Ac, Glycolysis

## Abstract

Nuclear paraspeckles assembly transcript 1 (NEAT1) is a well-known long noncoding RNA (LncRNA) with unclear mechanism in Alzheimer’s disease (AD) progression. Here, we found that NEAT1 down-regulates in the early stage of AD patients and APPswe/PS1dE9 mouse. Moreover, knockdown of NEAT1 induced de-polymerization of microtubule (MT) and axonal retraction of nerve cells by dysregulation of the FZD3/GSK3β/p-tau signaling pathway. Histone acetylation analysis at the Frizzled Class Receptor 3 (FZD3) promoter shows a marked decreased in the levels of the H3K27 acetylation (H3K27Ac) after NEAT1 knockdown. Our data demonstrates that P300/CBP recruited by NEAT1 to the FZD3 promoter and induced its transcription via histone acetylation. In recent years a growing number of evidences have shown an abnormal brain glucose homeostasis in AD. In the present study we also observed an abnormal brain glucose homeostasis and enhanced sirtuin1 (SIRT1) activity after knockdown of NEAT similarly as in AD. Our results provided insight into the role of NEAT1 in the maintenance of MT stability and its effect on glucose metabolism during early stages of AD.

## Introduction

Alzheimer’s disease (AD) is the leading cause of dementia among the aging population that involves complex neurodegenerative alterations. There are several hypotheses to explain the basis of AD. Among them, the cholinergic, amyloid-β (Aβ) and tau hypotheses are the most recognized doctrine [1–4]. Currently, the available therapy of enhancing the acetylcholine response is not much satisfactory, and the trials targeting Aβ in AD repeatedly failed [5]; therefore, the microtubule associated protein tau (MAPT) hypothesis has gained much attention. In animal model, hyper-phosphorylated tau induces neurofibrillary tangles and microtubule disintegration, triggering the death of neurons [6, 7]; but, the precise molecular mechanism leading to the hyper-phosphorylation of tau remains unclear.

Nuclear enriched abundant transcript 1(NEAT1) is a type of nuclear bodies exist in highly organized manner in mammalian nuclei to control gene expression and epigenetic events, is critical for the formation and maintenance of paraspeckles, [8, 9]. The NEAT1 gene has two isoforms, NEAT1v1 (3.7 kb in length) and NEAT1v2 (23 kb in length). NEAT1v2 binds directly to the paraspeckles proteins P54nrb/NONO and SFPQ/PSF, which results in the recruitment of NEAT1v1 and PSPC1 [10]. The primary function of paraspeckles is to sequester A-to-I hyper-edited RNAs which composed of an inverted repeat of Alu elements (IRAlus) in the nucleus to prevent the incorrect translation of edited RNAs. Chen and Carmichael investigated that, NEAT1 knockdown leads to the collapse of paraspeckles as well as increased export of mRNA containing IRAlus, suggesting a close relationship between NEAT1 and the function of nuclear retention[12]. In addition to its involvement in the regulation of multiple genes expression by modulating their transcriptional activities NEAT1 also implicated in some neuronal loss diseases and neurodegenerative processes, such as amyotrophic lateral sclerosis, traumatic brain injury and Huntington’s disease., However very little is known about the precise roles of NEAT1 in AD progression [13–15]. In our recent study, we found that NEAT1 expression down-regulated in the early stage of AD and its downregulation contribute to amyloid-β deposition [16] and the change of glucose metabolism.

Several lines of studies have shown that abnormal glucose homeostasis in the brain associates with AD pathogenesis. Increased in brain glucose levels and reduced in glycolytic influx due to impair glucose metabolism seems to be an intrinsic features of AD, appear early before the onset of clinical symptoms[17]. Epidemiological studies indicate that elevated blood glucose levels are a risk factor for dementia. Patients with type 2 diabetes are more likely to develop AD, suggesting that abnormal glucose metabolism plays an instrumental role in the development of AD [18, 19]. In this study, we found that NEAT1 down-regulation during early stage of AD disturbed glucose metabolism of neuron cells and impaired microtubule (MT) stabilization via dysregulation of the FZD3/GSK3β/p-tau signaling pathway. Metformin abandon tau hyper-phosphorylation and axonal degeneration via increased NEAT1 expression. Our results suggest new insights into the relationship between AD and diabetes mellitus.

## Results

### NEAT1 silencing induces de-polymerization of microtubules (MTs) during early stages of Alzheimer Disease (AD)

With the help of Kolmogorov-Smirnov test, we analyzed, the expression profiles in hippocampus of different stage AD patients and normal persons using data from the National Center for Biotechnology Information (NCBI). The expression levels of NEAT1 in the hippocampus of AD patients at different stages (GSE84422) is shown in figure 1A. NEAT1 expression significantly reduced in Braak stage 1 and 2, representing an early-stage of AD, compared to patients in other Braak stages or normal persons (Fig 1A). Braak stage 1 and 2 are the earliest disease phases in AD, in which abnormal tau and neurofibrillary tangle start to appear. Interestingly in our study, we observed a significantly reduced Neat1 expression levels in the hippocampus of 2-month-old APPswe/PS1dE9 and C57 mice (wild type) representing an early stage of AD (Fig 1B).

**Figure 1.**
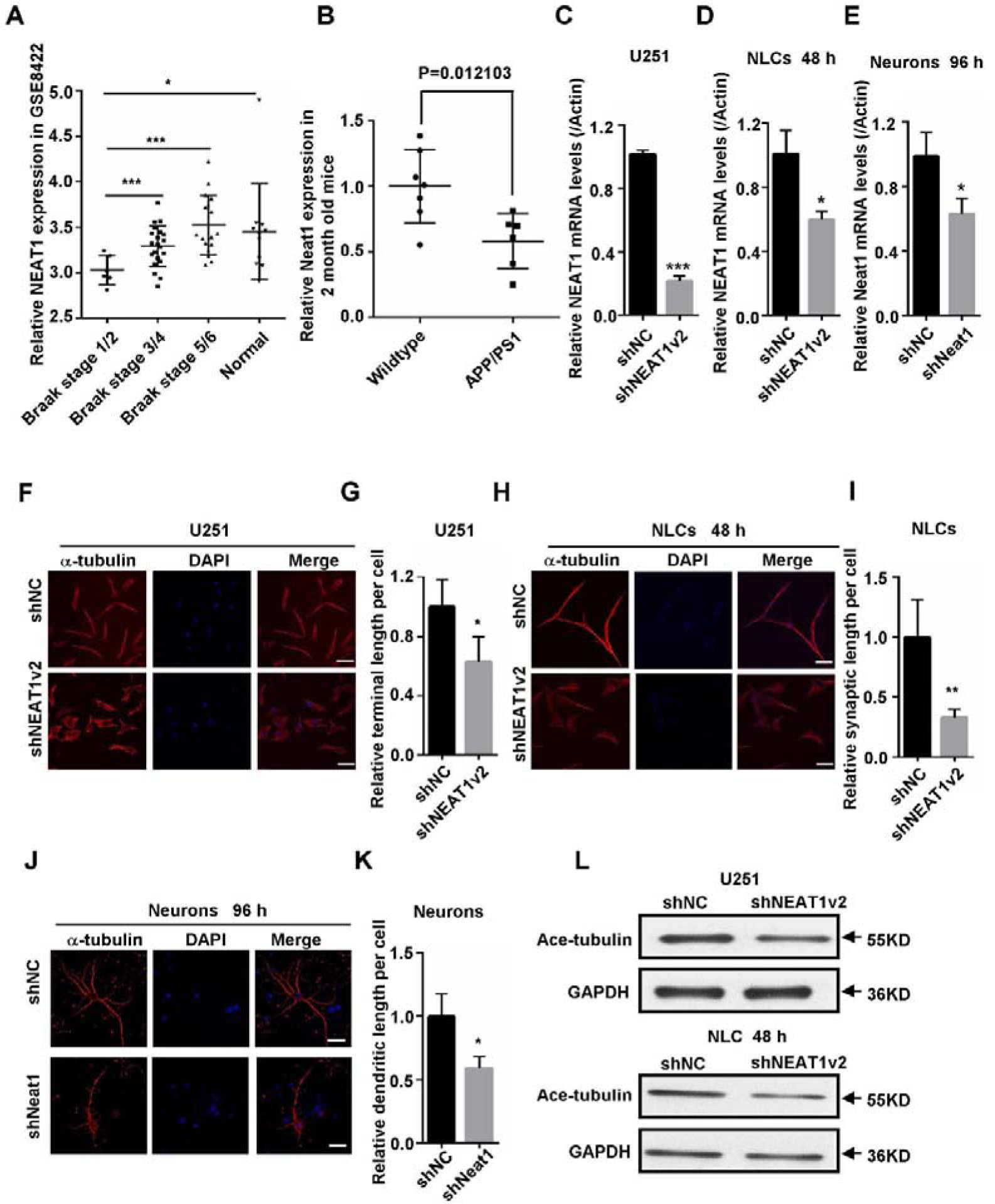
NEAT1 silencing induces de-polymerization of microtubules (MTs) during early stages of Alzheimer Disease (AD). A. The expression of NEAT1 in the hippocampus of AD patients with different braak stage and normal persons was analyzed in GSE84422. B. Neat1 analysis in hippocampi of 2-month-old AD mice and wild-type mice. C. The NEAT1 mRNA expression levels were measured by quantitative PCR in stable NEAT1 deficient cell line on U251. D. Quantitative PCR analysis of NEAT1 mRNA levels in lentivirus based shNEAT1v2 and shNC transfected NLCs. E. Neat1 mRNA expression was detected by quantitative PCR in lentivirus based shNeat1v2 and shNC transfected primary mouse neurons. F. Immunofluorescence analysis of α-tubulin (red) in stable NEAT1 deficient cell line on U251, H. shNEAT1v2 and shNC transfected NLCs, J. shNeat1v2 and shNC transfected mouse primary neurons. DAPI (blue) was used to stain the nuclei. Scale bars, 20μm. G. The average terminal length of stable U251 cell lines was calculated from 30 randomly selected cells. I. The average synaptic length of shNEAT1v2 and shNC transfected NLCs was calculated from 30 randomly selected cells. K. The average dentritic length of shNeat1 and shNC transfected mouse primary neurons was respectively calculated from 30 randomly selected cells. Image J software was used to count the cell length (mean ± s.d, \**P* < 0.05, \*\**P* < 0.01, \*\*\**P* < 0.001, Student 2-tailed *t* test). L. Western blot analysis for acetylated tubulin expression levels in stable U251 cell lines as well as shNEAT1v2 and shNC transfected NLCs.

To know the effect of down-regulation of NEAT1 on nerve cells, first we generated a NEAT1-deficient (shNEAT1v2) and negative control (shNC) stable cell lines on U251 using lentivirus based shRNA with inhibition efficiency approximately up to 80% (Fig 1C). After that, Mesenchymal stem cells (MSCs) from human placenta treated with suitable neuron-specific induction medium to induce neuron-like cells (NLCs), displayed neuron-protrusion morphologic changes during the first 4-6 days (Appendix Fig S1A). The expression of neuronal markers assessed by real-time PCR showed that GFAP (astrocyte marker), NF-M (neurofilament medium) and NSE (neuron-specific enolase) (neuronal marker) up-regulate in differentiated cells compared to the untreated MSCs (Appendix Fig S1B), [20–22]. The levels of NEAT1 mRNA declined significantly after transfection with lentivirus based shNEAT1v2 compared to shNC (small RNA with random sequence) in NLCs (Fig 1D). Moreover, the primary mouse neurons also transfected with lentivirus based shNeat1v2 and shNC, resulted in 50% inhibition ratio (Fig 1E).

We performed an Immunostaining with anti-α-tubulin antibodies in these cells and observed that down-regulation of NEAT1 induced the shrinkage of U251 cells (Fig 1F and Appendix Fig S1C) and significantly decreased the length of glial cell terminals in shNEAT1v2 cells compared to shNC cells (Fig 1G). Next we examined whether NEAT1 down-regulation have any effect on neurites extension of NLCs or mouse primary neurons. Immunofluorescence analysis (Fig 1H-K) and optical microscopic observation (Appendix Fig S1D), showed decreased in the axonal length and shrinkage of NEAT1 knockdown NLCs or primary mouse neurons, suggesting that depolymerization of MTs and degeneration of axons is mediated by the down-regulation of NEAT1, an important pathological changes towards AD development. The normal primary mouse neurons were also immunostained by α-tubulin antibody (Appendix Fig S1E). The western blot analysis shows the decreased expression of acetylated tubulin in NEAT1 deficient U251 cell line and NLCs, representing a reduction in polymerized tubulin[23], therefore suggesting an aggravated de-polymerization of MTs (Fig 1L)

### NEAT1 silencing mediates de-polymerization of MTs via FZD3/GSK3β/p-tau signaling pathway

To further investigate the potential mechanisms of NEAT1 in AD patients, a correlation analysis performed by using hippocampus samples. The expression profiles of (GSE84422) from GEO datasets, and a cluster (|r| > 0.4) of NEAT1-associated genes obtained from braak stage 1/2 AD patients. Moreover, GO analysis performed by using the NEAT1-positive correlation genes. Expression profile of GSE84422 showed that NEAT1 primarily associate with Wnt signaling pathway (Fig 2A). The results of GO analysis also showed that Wnt pathway may have some relationship with changes in NEAT1 expression during early stage of AD patients. We observed the expression of some important proteins in Wnt signaling pathway and found that the level of FZD3 mRNA in NEAT1 deficient U251 cell and NLCs reduced considerably compared to shNC (Fig 2B and C). We also found that siNEAT1v2 similarly to siFZD3 also down-regulated FZD3 expression in U251 (Fig 2D).

**Figure 2.**
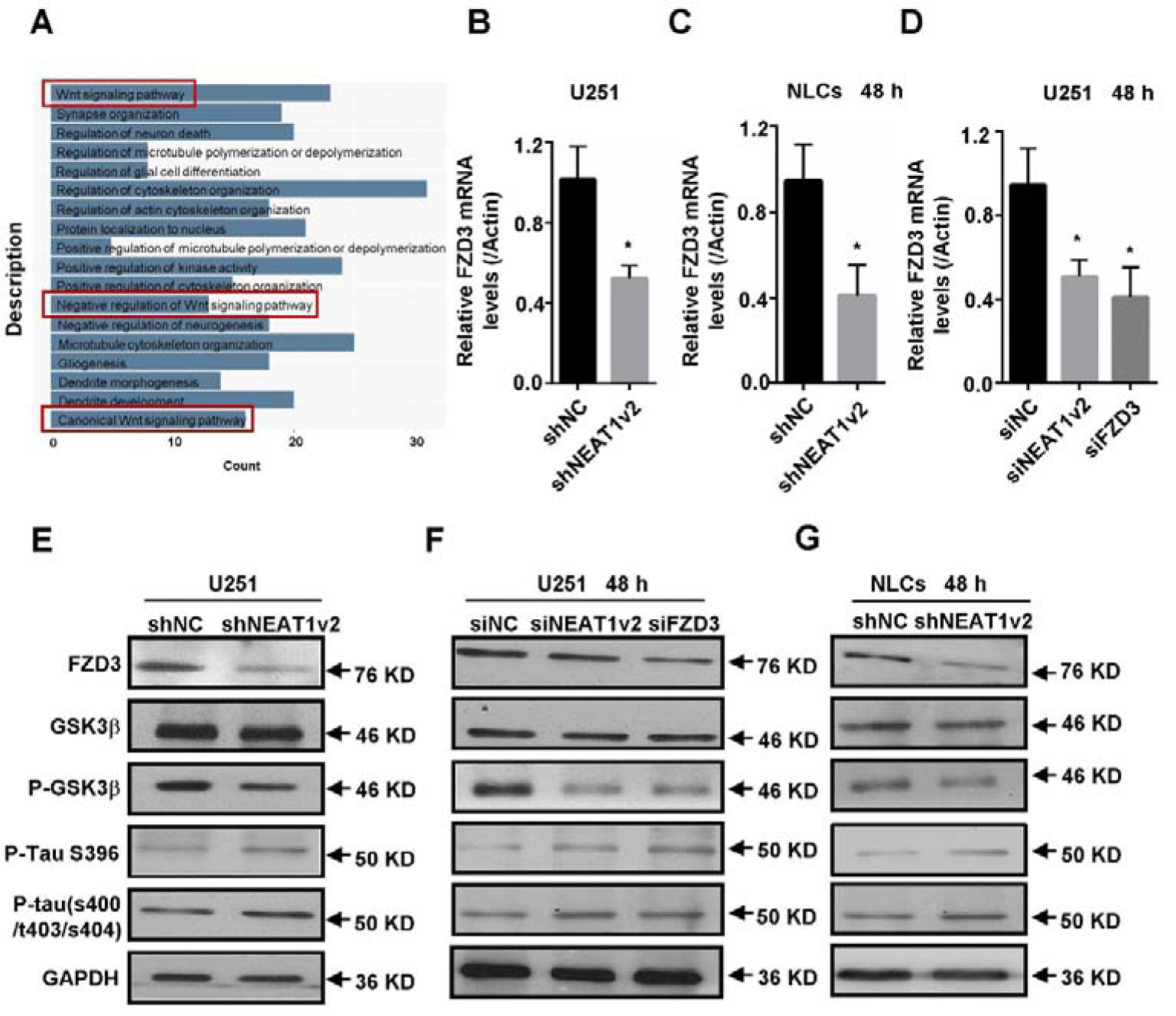
NEAT1 silencing mediates de-polymerization of MTs via FZD3/GSK3β/p-tau signaling pathway. A. GO analyses were performed using the NEAT1 positive-associated genes in GSE84422. B. The FZD3 mRNA expression levels were measured by quantitative PCR in stable NEAT1 deficient cell line on U251. C. The mRNA levels of FZD3 in lentivirus based shNEAT1v2 and shNC transfected NLCs. D. The mRNA levels of FZD3 in U251 after being transfected with NEAT1 siRNA, FZD3 siRNA and negative control. (mean ± s.d, \**P* < 0.05, Student 2-tailed *t* test). E-G. The expression levels of the FZD3, GSK3β, p-GSK3β, p-Tau(s396) and p-Tau(s400/t403/s404) were analyzed with Western blotting respectively in stable NEAT1 deficient cell line on U251, NEAT1v2 siRNA, FZD3 siRNA transiently transfected U251 and shNEAT1v2 and shNC transfected NLCs.

In addition to its role in Wnt signaling, FZD3 regulate GSK3β activity. Activation of disheveled protein 1(Dv1) by FZD3 suppress GSK3β activation via Dv1-mediated inhibition[24]. Glycogen synthase kinase 3β (GSK3β) is also an important tau protein kinase [4, 25, 26]. Therefore, we next examined the level of P-GSK3β, P-tau and FZD3 proteins in NEAT1 deficient U251 cell line, NEAT1 siRNA, FZD3 siRNA transfected U251 cells and shNEAT1v2 transfected NLCs. The results show that NEAT1 knockdown significantly decreased the expression of FZD3 and p-GSK3β (ser9) but the level of total GSK3β remain unchanged, suggesting a decreased in p-GSK3β/t-GSK3β ratio (i.e., increased GSK3β activity). Phosphorylation of GSK3β at serine 9 (p-GSK3β) is a modification process required to inhibits GSK3β activity [4, 25]. The decreased in p-GSK3β /t-GSK3β ratio thereby increased an active form of GSK3β, and ultimately result an elevated level of Phosphorylated form of Tau protein (Fig 2E-G).

### NEAT1 regulates FZD3 expression via interaction with P300/CBP

To understand the molecular mechanism of NEAT1-regulated FZD3 expression, we generated luciferase reporter constructs containing the promoter region of FZD3 and transfected the reporter vector into NEAT1 deficient U251 cell line. The results of the luciferase activity assay showed that NEAT1 knockdown reduced the transcriptional activity of the FZD3 promoter, indicating that NEAT1 regulate the expression of FZD3 at transcriptional level, (Fig 3A). H3K27Ac is a well-established marker for active transcription, we therefore analyzed its changes by using chromatin immunoprecipitation (ChIP) assays and the results reveal a broad and significantly decreased enrichment of H3K27Ac at the FZD3 promoter (Fig 3B).

**Figure 3.**
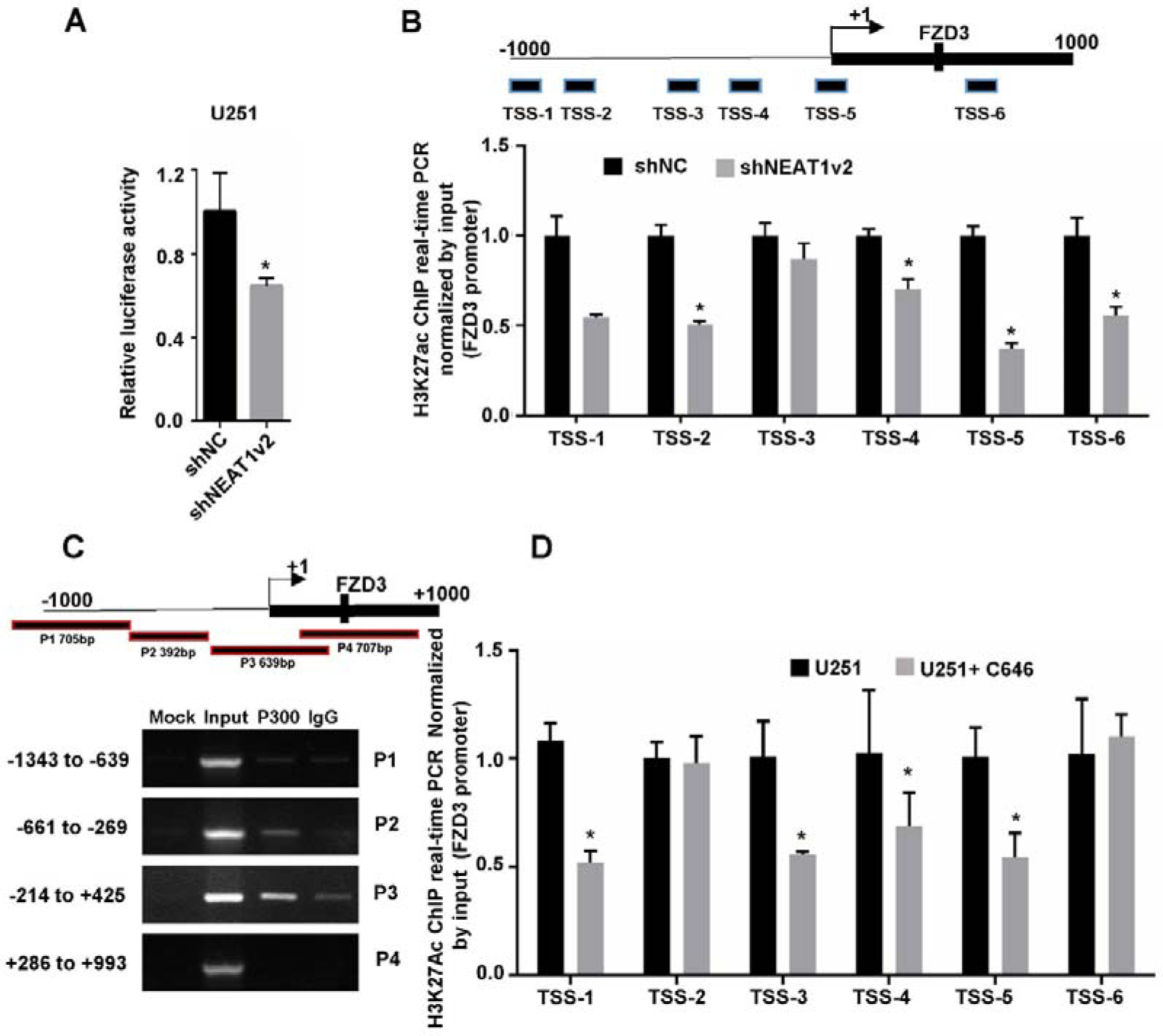
NEAT1 silencing decreases FZD3 expression via recruitment of P300/CBP and reduced level of H3K27Ac. **A.** After co-transfection with siNEAT1v2 or negative control siRNAs and the pGL3 enhancer plasmid containing FZD3 promoter fragments for 48h, the relative transcriptional activities were determined with a luciferase assay in three independent experiments. **B.** The histone modification status of H3K27Ac on FZD3 promoter detected by ChIP assays in stable NEAT1 deficient cell line on U251. D. ChIP-PCR showed that P300/CBP occupied the FZD3 promoter region. D. The 20μM C646 treated and normal U251 cells were collected for ChIP assays to analyse the relative fold enrichment of the FZD3 promoter using anti-H3K27Ac antibody. (mean ± s.d, \**P* < 0.05, \*\**P* < 0.01, Student 2-tailed *t* test).

Genome-wide profiling analysis using ChIP-sequencing have identified thousands of functional/active enhancers that are either bound by the transcriptional co-activator p300/CBP, or characterized by their association with high levels of H3K27Ac[27]. In our previous study of RNA immunoprecipitation (RIP) assay we have shown that p300/CBP pulled down several fragments of NEAT1. Moreover NEAT1 also co-localized with p300/CBP therefore influence its acyltransferase activity by direct interaction[16]. To reveal the potential mechanism how NEAT1 regulates FZD3 transcription, we designed four paired primers (P1, P2, P3, P4) across the promoter region (−1000bp to +1000bp) of FZD3 and then subjected to ChIP-PCR assay to identify its regulatory mechanism. The results showed that P300/CBP likely to bind to P2 region of FZD3 promoter (Fig 3C), which is -661bp to -269bp from transcription start site of FZD3. To further clarify the relationship between p300/CBP and FZD3, we cultured normal U251 with 20 uM C646, a selective inhibitor of p300, for 48h and observed a significant reduction of H3K27Ac enrichment at FZD3 promoter compared to untreated U251(Fig 3D). Taken together, our preliminary results suggest that NEAT1 recruits p300/CBP at FZD3 promoter region therefore influence its transcription due to change in H3K27Ac levels.

H3K4Me3 is an active transcription marker, whereas H3K27Me3 suppress transcription. We therefore examined the methylation pattern of histones proteins including tri-methylated histone H3 at lysine 4 (H3K4Me3) and tri-methylated histone H3 at lysine 27 (H3K27Me3), after down-regulation of NEAT1. The results show that decreased enrichment of H3K4Me3 and increased enrichment of H3K27Me3 at the FZD3 promoter region, is due to the down-regulation of FZD3 (Appendix Fig S2A and B) in NEAT1 down-regulated U251 stable cell line. Overall, our data suggest that NEAT1 knockdown enhance the phosphorylation of tau by inhibiting FZD3 transcription whereas activating the GSK3β signaling pathway (Appendix Fig S2C).

### NEAT1 silencing decreases FZD3 expression via impaired glycolysis and increased SIRT1 activity

During our investigation, we observed that in similar cell numbers (Appendix Fig 3A and B), NEAT1 deficient U251 cells produced less acid in medium compared to control cells as evident of slow change in color to yellowish (Fig 4A), suggesting that knockdown of NEAT1 induced down-regulation of glycolysis and its acidic metabolites. To further confirm the glycolysis of NEAT1 deficient U251 cells, we used Seahorse ECAR (Extracellular Acidification Rate). In Figures 4B and C, glycolysis and the glycolytic capacity shows a 46.5% and 40.1% decreased respectively in NEAT1 stable knockdown cells (Fig 4B and C). Consistent with the color change of the culture medium, the intracellular pH value detected by flow cytometer also increased in shNEAT1v2 cells (Fig 4D). NEAT1 silencing also caused a decline in the expression profile of multi-glycolysis related genes, including PFKFB2, PKM2 and HK2 (Fig 4E). Furthermore, NEAT1 silencing-mediated down-regulation of glycolysis increased the ratio of NAD/NADH and the activity of SIRT1 (Fig 4F and G).

**Figure 4.**
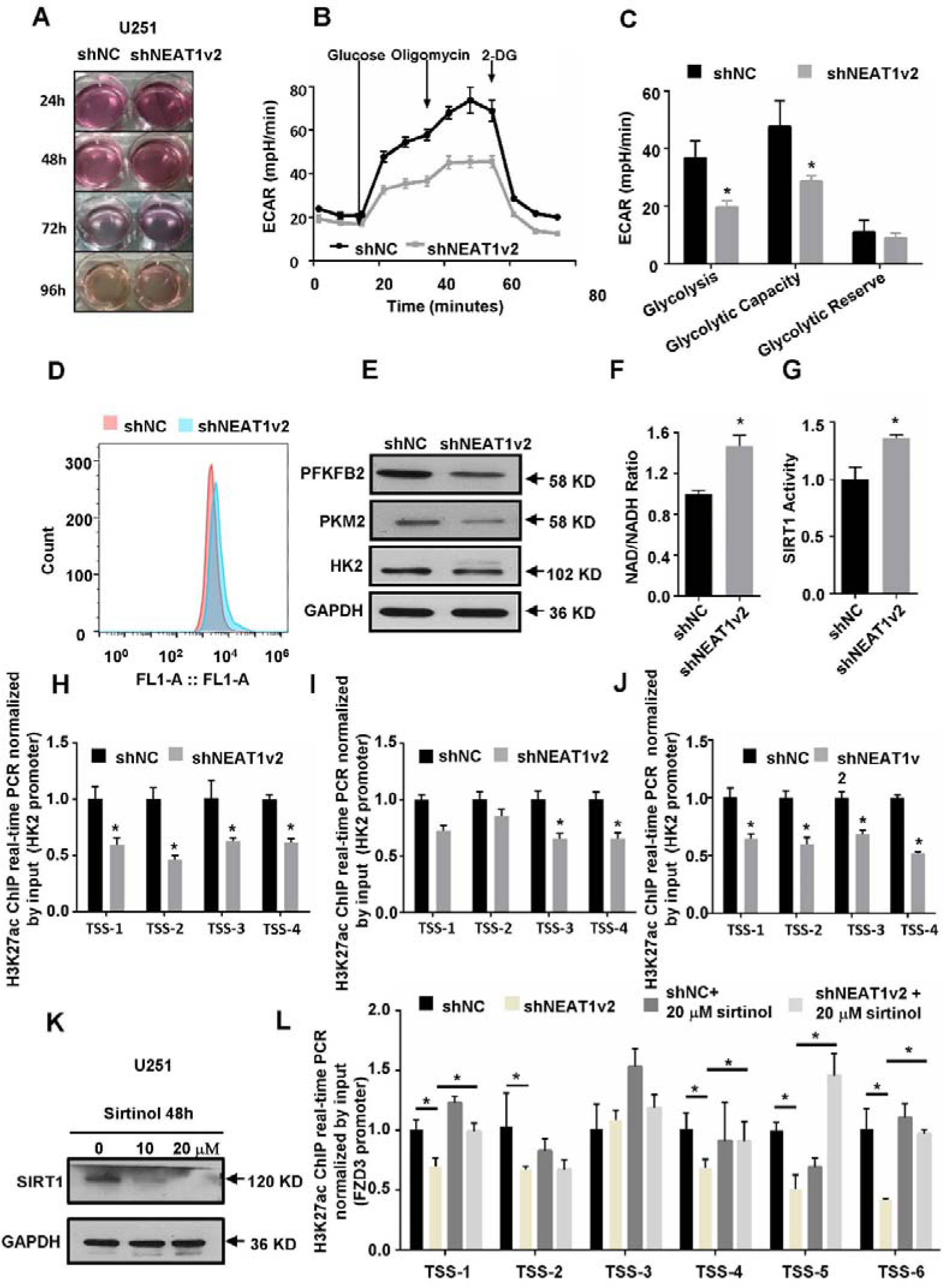
NEAT1 silencing decreases FZD3 expression via impaired glycolysis and increased SIRT1 activity. **A.** The color change of stable NEAT1 deficient cell lines were cultured in a 6-well plate for different time. B. The Seahorse Flux Bioanalyser was used to measure ECAR of stable NEAT1 deficient cell lines in real time under basal conditions and in response to glucose, oligomycin, and 2-Deoxy-D-glucose (2DG). n=3, *P < 0.05. C. Basal glycolysis (after the addition of glucose), glycolytic capacity (after the addition of oligomycin), and glycolytic reserve (calculated as the difference of oligomycin rate and glucose rate) ECAR. n=3, *P < 0.05. D. The value of intracellular pH in stable NEAT1 deficient cell lines detected with Flow cytometry by using fluorescent dye, BCECF. E. Immunoblotting analysis of PFKFB2, PKM2, and HK2 in stable NEAT1 deficient cell lines on U251. F. Analysis of NAD+ / NADH ratio in stable NEAT1 deficient cell lines on U251. G. The assay of SIRT1 activity in stable NEAT1 deficient cell lines on U251. H-J. ChIP assays of H3K27Ac enrichment on the promoters of HK2, PFKFB2 and PKM2, the fold enrichment of H3K27Ac on these promoter fragments was standardized relative to the input level. K. Western blotting analysis of SIRT1 expression after treated with 0, 10, 20μM Sirtinol for 24h. L. Using ChIP analysis to detect the H3K27Ac enrichment on FZD3 promoter regions when the NEAT1 deficient cell lines were treated with 20 μM Sirtinol for 48h. \**P* < 0.05, \*\**P* < 0.01, Student 2-tailed *t* test.

To investigate how NEAT1 regulate cellular glycolysis, we detected H3K27Ac enrichment on the promoter regions of these glycolysis related genes. The results showed that NEAT1 knockdown significantly reduced H3K27Ac enrichment levels when compared with shNC cell line (Fig 4H-J). SIRT1 is a nuclear enzyme of NAD-dependent histone deacetylase, and its catalytic activity mainly depends on NAD/NADH. An increased in the ratio of NAD/NADH may attenuates H3K27Ac enrichment at the promoter regions of FZD3 or other genes, due to enhanced activity of SIRT. We therefore sought to know whether increased in SIRT1 activity is mediated by NEAT1 silencing which contributes to FZD3 transcription, SIRT1 inhibitor Sirtinol added to cultured shNEATv2 deficient cells to analyze alterations of H3K27Ac enrichment at FZD3 promoter region. The results reveal a significantly increased enrichment of H3K27Ac at the FZD3 promoter after adding 20μM Sirtinol for 48 h, suggesting an increased in SIRT1 activity to some level therefore may influence FZD3 transcription (Fig 4K and L).

### Metformin abandon tau hyper-phosphorylation and axonal degeneration via increased NEAT1 expression

Metformin originally been used as an anti-diabetic agent, but recently, it is considered as promising agent for the treatment of AD [28]. We therefore investigated whether metformin can retard tau hyper-phosphorylation and axonal degeneration. To apply an optimum concentration of metformin, first we calculated LC50 of the drug (from 0 to 15mM) for U251 cells by using cell counting kit 8 assay. The results show that cell viability not influenced significantly by low concentration (from 0 to 2mM) of metformin (Appendix Fig 3C). We also observed that metformin concentration dependently increased NEAT1 as well as FZD3 expression in U251 cells. Metformin concentration ranged from 0 to 0.5 or 1mM increased the expression of NEAT1 or FZD3 and reached to the maximum level after that decreased steadily (Fig 5A).

**Figure 5.**
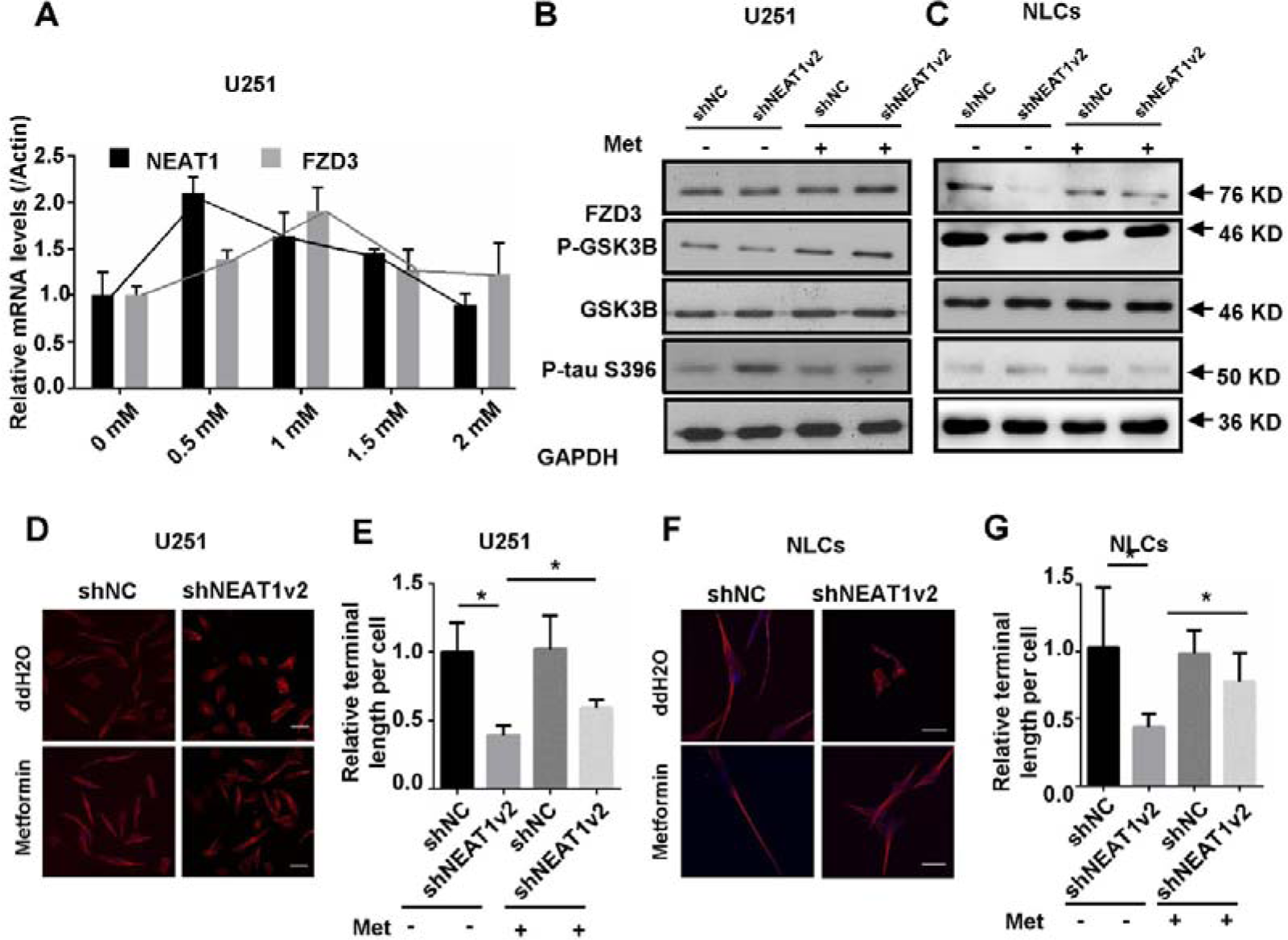
Metformin abandon tau hyper-phosphorylation and axonal degeneration via increased NEAT1 expression. A. Quantitative PCR analysis of NEAT1 and FZD3 expression of 0 to 2mM metformin treatment on normal U251 cells. B. Expression levels of FZD3, P-GSK3β, GSK3β, P-tau(ser396) was detected by Western blotting after 48h treatment of 0.5mM metformin in stable NEAT1 deficient cell lines on U251. C. Expression levels of FZD3, P-GSK3β, GSK3β, P-tau(ser396) was detected by Western blotting after 48h treatment of 1mM metformin in lentivirus based shNEAT1v2 and shNC transfected NLCs. D. Metformin treated stable NEAT1 deficient cell lines on U251 were stained with antibody against α-tubulin(red) and subjected to confocal microscopy analysis. DAPI(blue) was used to stain the nuclei. Scale bars 20 μm. E. B. The average terminal length of stable NEAT1 deficient cell lines treated with metformin was calculated from 30 randomly selected cells, respectively. F. Metformin treated shNEAT1v2 and shNC transfected NLCs were stained with antibody against α-tubulin(red) and subjected to confocal microscopy analysis. DAPI(blue) was used to stain the nuclei. Scale bars 20 μm. G. The average synaptic length of shNEAT1v2 and shNC transfected NLCs treated with metformin was calculated from 30 randomly selected cells. Image J software was used to count the cell length. (mean ± s.d, \**P* < 0.05, Student 2-tailed *t* test).

The expression levels of downstream genes, including FZD3, P-GSK3β, as well as tau hyper-phosphorylation also rescued by 0.5 mM metformin in NEAT1 deficient U251 cells (Fig 5B). Furthermore, expression of FZD3, P-GSK3β and tau hyper-phosphorylation also reduced by 1mM metformin in shNEAT1v2 and shNC transfected NLCs (Fig 5C). We also observed that compared to control, treatment with metformin, rescued the glial cell terminal length in U251 cells and the average terminal length analyzed by Image J (Fig 5D and E). Interestingly the rescued axonal length also observed in shNEAT1v2 and shNC transfected NLCs (Fig 5F and G and Appendix Fig S3D). However, high concentration of metformin not only failed to rescue the axonal degeneration but further triggered cell lesion and even death in NLCs, showing its cytotoxic effect in neuronal cells (Appendix Fig S3E).

### Metformin abandon tau hyper-phosphorylation in hippocampus of APPswe/PS1dE9 double transgenic mice

We next examined Neat1 expression in the hippocampus of 2-month-old wild type and AD transgenic mice, showing an early stage of AD (Fig 1B). We also examined Fzd3 expression levels in 2-month-old mice and found decreased in mRNA levels of Fzd3 in hippocampus of AD mice (Fig 6A). The rescuing effects of metformin further confirmed by in-vivo studies. In our previous study, we detected the mRNA levels of Neat1 in 3-month-old APP/PS1 double transgenic mouse, and found reduced Neat1 expression compared with C57 mice [16]. In present study, these nearly 3 months old AD mice were observed an increased in the level of Neat1 and Fzd3 mRNA in the hippocampus of metformin treated group compared to control group treated with ddH2O (Fig 6B and C). Furthermore, we found increased H3K27Ac protein expression and reduced P-tau level in metformin treated AD mice group compare with ddH2O group (Fig 6D).

**Figure 6.**
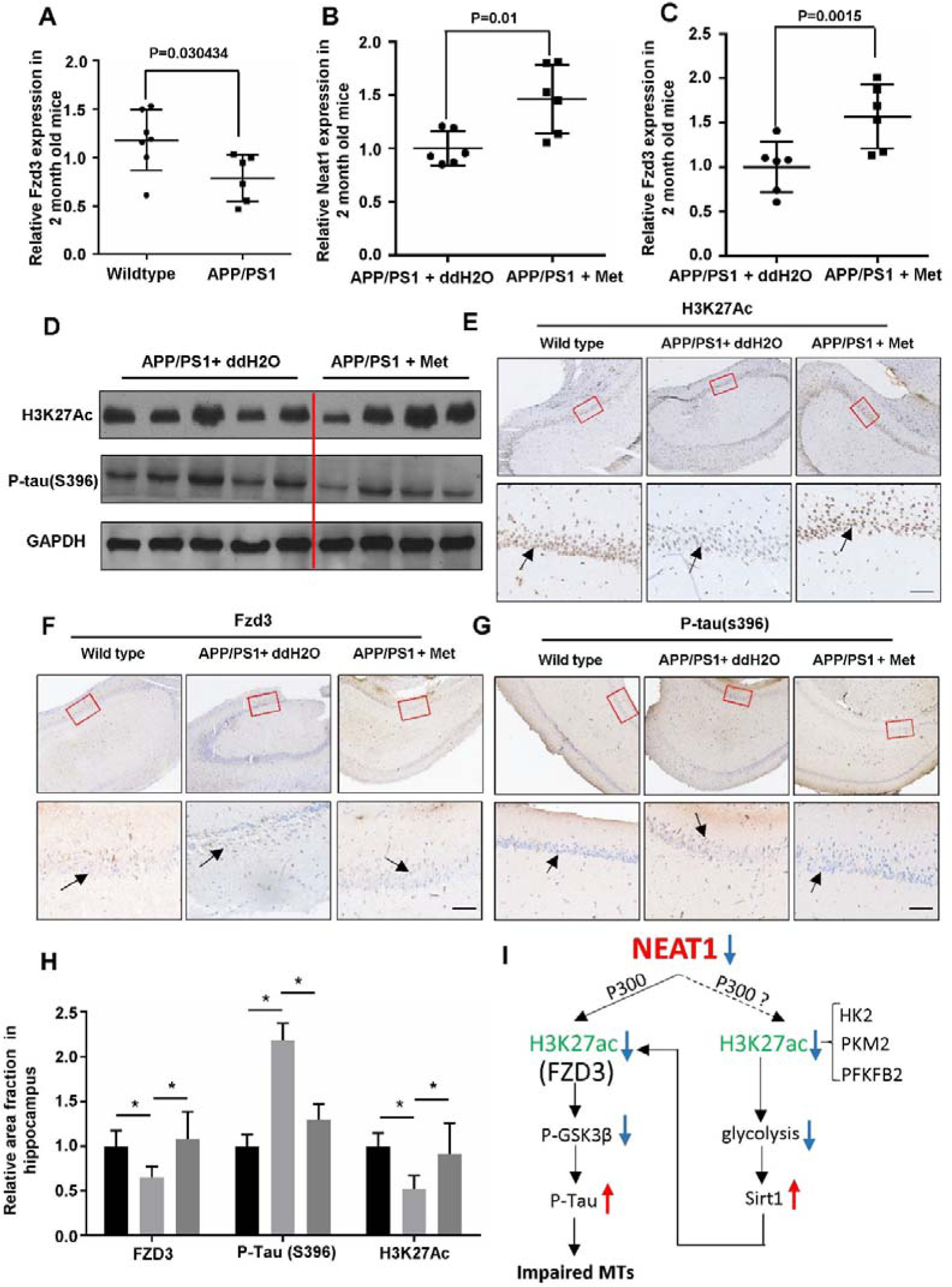
Metformin abandon tau hyper-phosphorylation in hippocampus of APPswe/PS1dE9 double transgenic mice. A. Fzd3 analysis in the hippocampi of 2-month-old APPswe/PS1dE9 double transgenic mice (n=6) and wild-type mice (n=7). B and C. Quantitative PCR analysis of Neat1 and Fzd3 expression of 4-weeks ddH2O treated (n=6) and metformin treated (n=6) APPswe/PS1dE9 double transgenic mice. D. Expression levels of P-tau(S396) and H3K27Ac were detected in the hippocampus of ddH2O and metformin treated APPswe/PS1dE9 double transgenic mice. E-G. The hippocampus of different treated mice was immunohistochemically stained with H3K27Ac, FZD3 and Phosphorylated Tau(Ser396), Scale bars 50μm. H. Quantification of the relative area fraction occupied by immunostaining of H3K27Ac, FZD3 and Phosphorylated Tau(Ser396) in CA1 of hippocampus were analyzed by Image J. (mean ± s.d, *P < 0.05, Student 2-tailed *t* test). I. Summary of the mechanism of downregulated NEAT1 during the early stage of AD

Next we performed Immunohistochemistry (IHC) in the whole hippocampus and in the CA1 hippocampal area of differently treated mice. The results show a significantly high H3K27Ac-positive and FZD3-positive neuronal cells in CA1 hippocampal area of metformin treated AD mice, compared to APPswe/PS1dE9 double transgenic mice treated with ddH2O. The accumulation of hyper-phosphorylated form of tau decreased significantly in the hippocampus, showing the rescue effect of metformin (Fig 6E-G). Quantification of the relative area fraction occupied by immunohistochemical staining was analyzed with Image J (Fig 6H).

## Discussion

NEAT1 is implicated in many neurodegenerative disorders, including amyotrophic lateral sclerosis (ALS) [13] and Huntington’s disease (HD) [9, 15]. In our recent study, we found that NEAT1 also involves AD progression. The down-regulation of NEAT1 expression contributed to amyloid-β deposition, via inhibiting expression of multi endocytosis related genes. Through interaction with P300/CBP complex, NEAT1 regulated expression of these genes via affecting H3K27 acetylation (H3K27Ac) and H3K27 crotonylation (H3K27Cro) in the promotor region of these genes[16]. In this investigation, we found that NEAT1 involves AD progression also through dysregulation of glycolysis and up-regulation of p-tau.

According to the early reports from other authors, the disorder of glucose metabolism has some relationship with AD. FDG PET study revealed that decrease utilization of glucose in the brain of patients with “sporadic” AD caused by gene mutation even before any visible symptoms of dementia, suggesting that disturbance in glucose metabolism is closely linked to AD development [29, 30]. In vivo diabetic model of mice demonstrates morphological and molecular alterations in the brain similar to those observed in other neurodegenerative disorders [31, 32]. However, the mechanism is still poorly understood. In our investigation, we found that NEAT1 expression markedly declines in the early-stage of AD patients. Down-regulation of NEAT1 mediates decrease in glycolysis and enhance the NAD/NADH ratio and the activity of SIRT1, which is a nuclear enzyme of NAD-dependent histone deacetylases (Fig 6I).

In addition to its function as a fundamental structural component of paraspeckles to regulate RNA nuclear retention and RNA splicing, NEAT1 involves in the epigenetic regulation of gene expression. Chen et al. reported that NEAT1 mediates H3K27me3 of Axin2, ICAT, and GSK3β in glioblastoma by binding to EZH2[33]. NEAT1 also associates with H3K4me3 [34, 35]. In the present study, we reported that NEAT1 regulates the tri-methylation and acetylation of H3K27 of the FZD3 promoter. Our investigation for the first time highlights NEAT1-mediated regulation of H3K27 acetylation at promoter region of the FZD3 gene. Through regulation of H3K27 acetylation of FZD3, NEAT1 activates GSK3β.

GSK3β play roles in both diabetes and AD progression. Insulin resistance dysregulates the insulin/PI3K/Akt/GSK3β signaling pathway, leading to the inhibition of Akt and de-phosphorylation of GSK3β (activation), result in tau hyper-phosphorylation [36–39]. The NEAT1/FZD3/GSK3β signaling pathway reported in this investigation also associate with hyper-phosphorylation of tau. Therefore, GSK3β appears to be a possible connection between the NEAT1/FZD3/GSK3β and insulin/PI3K/Akt/GSK3β pathways in AD progression. Our result also suggests that increased in SIRT1 activity mediated by NEAT1 down-regulation at least partially contribute to the change of enrichment of H3K27Ac at the FZD3 promoter region and the activity of FZD3/GSK3β signaling pathway (Fig 6I).

Metformin has been recommended as a first-line therapy for patients with type 2 diabetes mellitus since 2009 [40]. Because it can penetrate the blood-brain barrier, metformin is also used as a promising drug for treating AD. Chen et al. reported that metformin improve memory impairment and inhibit neuronal apoptosis and accumulation of Aβ in the hippocampus of db/db mice [41]. In our study, we reported that metformin rescued NEAT1 down-regulation and axonal degeneration by regulating the FZD3/GSK3β/p-tau signaling pathway. Consistent with our finding, researchers also reported that both in vitro and in vivo, metformin reduce tau phosphorylation in murine neurons, but via the mTOR/protein phosphatase 2A (PP2A) signaling pathway [42]. However, Barini et al., reported that metformin treatment in a mouse model of tauopathy promotes tau aggregation and exacerbates abnormal behavior[28]. Moore et al. reported that using metformin aggravated cognitive impairment of AD patients compared to control group. They think that MET-induced vitamin B12 deficiency may cause AD[43]. In our study, we found that high dose of metformin is cytotoxic for neuron cells. In short, when metformin is use to prevent or treat AD, dose is an important consideration.

### Experimental Procedures

#### Mice

Animals were kept in an environmentally controlled breeding room (temperature: 20 ± 2 °C; humidity: 60% ± 5%; 12 h dark/light cycle). The animals were fed standard laboratory chow diets with water ad libitum. The study was performed in strict accordance with the recommendations of the Guide for the Care and Use of Laboratory Animals of the Institutional Animal Care and Use Committee of Tsinghua University. The protocol was approved by the Animal Welfare and Ethics Committee of Tsinghua University, China. C57BL/6 (wild type mice) and APPswe/PS1dE9 double transgenic mice (transgenic AD mice) both at the ages of 2 months were obtained from Jackson Laboratory (Bar Harbor, ME, USA) and were housed in individually ventilated cages. 2 month old C57BL/6 (n=7) and same age APPswe/PS1dE9 double transgenic mice (n=6) were killed by decapitation to obtain hippocampus tissue and detected mRNA levels of Neat1 and Fzd3.

2 month old C57BL/6 (n=8) and same age APPswe/PS1dE9 double transgenic mice (n=12) were obtained from the same source and were divided into 3 groups: 1) a sham group (which only received normal diet, daily intragastric administration of ddH2O, C57BL/6 (n=8).) 2) a control group (which only received normal diet, daily intragastric administration of ddH2O, transgenic AD mice (n=6).) 3) a case group (which received daily intragastric administration of 200mg/kg metformin (Abcam, #ab120847) dissolved in ddH2O for 9 weeks, transgenic AD mice (n=6).) For Immunostaining, WT mice and AD mice were killed by decapitation to obtain hippocampus tissue and perfusion were performed with 4% PFA in PBS. To quantify Neat1 expression levels, C57BL/6 mice and AD mice were anesthetized with pentobarbital and hippocampal tissue was eluted in RNAiso Plus (Takara, D9108B) followed by RNA extraction.

#### Dataset

MRNA expression data sets and the associated clinical information were obtained from GSE84422 (GEO, https://www.ncbi.nlm.nih.gov/gds/) database. GSE84422 is titled as molecular signatures underlying selective regional vulnerability to Alzheimer’s Disease, which includes RNA samples from 19 brain region isolated from the 125 specimens. NEAT1 expression in the hippocampus of 44 AD patients with different braak stage and 11 normal persons was analyzed using nonparametric Kolmogorov-Smirnov test.

#### Cell culture

Human glioma U251 cells (China Infrastructure of Cell Line Resources) were cultured in Dulbecco’s modified Eagle’s medium (Gibco/Invitrogen Ltd, 12800-017) containing 10% fetal bovine serum (PAA, A15-101), 10 U/ml penicillin-streptomycin (Gibco/Invitrogen Ltd, 15140-122) in a 5% CO2-humidified incubator at 37 °C.

Human placenta derived mesenchymal stem cells (MSCs) were isolated and cultivated according to a previously published protocol [44]. MSCs (5×10^4^ per well on a 6-well culture plate) were cultured in low-glucose Dulbecco’s modified Eagle’s medium (LG-DMEM) containing 10% fetal bovine serum (FBS), 10U/mL penicillin/streptomycin at 37°C, and 5% (v/v) CO_2_. Primary hippocampal neurons were isolated from embryonic E18.5 C57BL/6 mice and approximately 2×10^3^ cells/well were plated on poly-D-lysine-coated glass coverslips (20 µg/ml) for immunofluorescence. The plating medium was DMEM-F12 (Gibco/Invitrogen Ltd, 10565-018) supplemented with 10% horse serum (Gibco/Invitrogen Ltd, 26050-070), 10mM sodium pyruvate (P4562, sigma), 0.5mM glutamine (G6392, sigma) and 1% D-Glucose (G6152, sigma). After 2-4 h, the medium was changed to neuronal growth medium (500ml Neurobasal medium (Gibco/Invitrogen Ltd, 21103-049) with 10ml B27(Gibco/Invitrogen Ltd, 17504-001), 5ml N-2 Supplement (Gibco/Invitrogen Ltd, A13707-01)10ul 45% D-Glucose (G6152, sigma) and 5ml L-Glutamine (Gibco/Invitrogen Ltd, A2916801)). 5ug/ml AraC (C6645, sigma) was added to neuronal growth medium after 72h to reduce glial growth. Lentivirus transfections were conducted as follows.

#### Cell transfections

All the synthetic lentivirus-based shRNAs (shNEAT1v2 and shNC) and siRNAs (siNEAT1v2 and siNC) were purchased from Shanghai GenePharma Co., Ltd. All the siRNAs were transfected with lipofectamine™ 2000 (Invitrogen, 11668-019) according to the manufacturer’s protocol, and shRNAs were co-transfected with polybrene (GenePharma Co. (Shanghai, China)). The siRNA sequences were showed in table S1.

#### Construction of stable cell lines

Human glioma U251 cells were cultivated in 6-well plate (1×10^4^/well). When the cell density reached 40-50% confluence, the Lentivirus based shRNAs were used for cell transfection to generate stable monoclonal cell lines (shNEAT1v2 cells and shNC cells). Puromycin (A1113803, Invitrogen; Thermo Fisher Scientific, Lnc) selection (10ug/ml) started 24h after transfection. The medium with 10 µg/ml puromycin was changed every 2-3 days. Following 2-4 weeks, isolated colonies were selected and grown for later assays.

### MSC induction

MSCs were incubated for 7-10 days in neuronal differentiation medium containing 10 ng/ml epidermal growth factor (EGF, sigma), 10 ng/ml basic fibroblast growth factor (bFGF, sigma), 10 µM Forskolin (Macklin, F823536, Shanghai), 10 ng/ml brain-derived neurotrophic factor (BDNF, sigma), 0.1 mM 3-isobutylmethyl-xanthine, IBMX (Macklin, 1811775, Shanghai), 0.5 µM retinoic acid (RA, sigma), 5% FBS and the DMEM/F-12, GlutaMAX™ medium (Dulbecco’s Modified Eagle Medium/Nutrient Mixture F-12, 10565018 Gibico). Using real-time PCR detects the expression changes of several neuronal marker genes after this induction, including GFAP, NF-M and NSE [45–47]. The primers were showed in table S2.

### Reverse Transcription and Quantitative PCR

Reverse transcription was performed using ReverTra Ace® qPCR RT Master Mix with gDNA remover (TOYOBO, FSQ-301) according to the manufacturer’s protocol. The resultant cDNA was measured by quantitative PCR using following system: 4µl of RNase-free H2O, 0.5 µl of forward primer (1 µM), 0.5ul of reverse primer (1 µM), 1 µl of cDNA (50 ng) template, and 5 µl of SYBR Green PCR Master Mix (TOYOBO, QPK-201) at 95°C for 30 s, followed by 40 cycles of 95°C for 15 s, 60°C for 15 s and 70°C for 15 s. All mRNA levels were normalized to beta-actin. The primers were showed in table S2.

### CCK8 assay

Approximately 5×10^3^ cells per well were plated in a 96-well culture plate and incubated 24 h, followed by treating different concentrations of metformin for 48 h. Then, 10 µl CCK8 reagent (MedChem Express, HY-K0301-500T, China) were added to each well and maintained for 4h at 37°CAfter that, the absorbance at 450nm was detected by using a microplate reader.

### Western blot

The antibodies used for Western blotting included an anti-GSK3β antibody (cell signaling, #12456), an anti-Phospho-GSK3β(ser9) antibody (cell signaling, #9322), an anti-FZD3 antibody (Abcam, ab75233), an anti-phospho-Tau (S396) antibody (Abcam, ab109390), an anti-Phospho-Tau (Ser400/Thr403/Ser404) antibody (Cell signaling, #11837), an anti-acetyl-Tubulin antibody (Sigma, T6793), an anti-PKM2 antibody (Cell signaling, #4053) an anti-HK2 antibody (Cell signaling, #2867), an anti-PFKFB2 antibody (Cell signaling, #13045) and an anti-GAPDH antibody (Proteintech,10494-1-AP) was analyzed by western blot. Protein sample was lysed in ice-cold whole cell extract buffer B (50 mM TRIS-HCl, pH 8.0, 4M urea and 1% Triton X-100), followed by being heated at 100°C for 10 min with 5× loading buffer. Then, equal amount of protein sample was separated by SDS-PAGE and transferred onto PVDF membranes (Millipore, Immobilon-NC). Membranes were blocked for 1h at room temperature with 5% non-fat milk and successively incubated overnight at 4°C with primary antibody and 2h room temperature for secondary antibody. After that, ECL Blotting Detection Reagents were used to visualize protein bands.

### Immunohistochemistry

The hippocampus of mouse was isolated and fixed in 4% paraformaldehyde (PFA) and then placed in 30% sucrose in PBS for 1 day. Paraffin-embedded tissue sections (10um thickness) were treated in 0.01M PBS containing 3% hydrogen peroxide (H_2_O_2_) for 10 min. Then blocked in 3% BSA and were incubated in following antibodies: an anti-phospho-Tau (S396) antibody (Abcam, ab109390), an anti-FZD3 antibody (Abcam, ab75233) and an anti-acetyllysine antibody (Cat#PTM-102, clone Kac-11). Specimens were visualized under an inverted phase contract fluorescent microscope [48].

### Immunofluorescence

Antibodies against α-tubulin (Sigma, T6074) and the secondary antibodies Alexa Fluor 594 (Life Technologies Corp.) were employed in immunofluorescence staining. Microscopic analysis that clearly reflecting changes of axonal length were captured using an Olympus FV1000 confocal laser microscope.

### Luciferase Assay

The dual-luciferase promoter assay system was generated by inserting sequences from −500bp to +500bp relative to the transcription start sites (TSS) of FZD3. The inserted reporters were obtained from Shanghai GenePharma Co. Ltd. Luciferase activities were assayed using a Dual-Luciferase Reporter System (Promega, E1960).

### ChIP Assay

ChIP assays were performed as described previously [49]. Briefly, the cells were digested with ChIP lysis buffer (50 mM Tris-HCL PH=8.0, 5 mM EDTA, 0.1% deoxycholate, 1% Triton X-100, 150 mM NACL in 1* PIC (protease inhibitor)), And were crosslinked with 1% formaldehyde and sonicated for 180s (10s on and 10s off) on ice shear the DNA to an average fragment size of 200-1000bp. The 500ul of sonicated chromatin was purified by centrifugation, and then, the supernatants were incubated with the ChIP grade antibody against 2-5ug anti-KAT3B/P300 (Abcam, ab54984), anti-Histone H3 (acetyl K27) (Abcam, ab4729), anti-Histone H3 (tri methyl K4) (Abcam, ab8580), anti-Histone H3 (tri methyl K27) (Abcam, ab6002) and 100ul Dynabeads^TM^ protein G (Invitrogen, 10004D, USA). Finally, chromatin DNA was subjected to Quantitative PCR and all primers for ChIP-qPCR are listed in table S3.

### NAD/NADH and SIRT1 deacetylase activity

The experiment was performed using NAD+/NADH Assay kit (Abcam, ab176723) according to the manufacturer’s instruction. SIRT deacetylase activity was measured using a colorimetric assay kit (Abcam, ab156915) and nuclear extracts were prepared using Nucleoprotein Extraction Kit (Sangon Biotech, C500009, Shanghai).

### Seahorse extracellular flux analysis

The extracellular acidification rate (ECAR), indicating glycolysis, was measured using Seahorse XFe96 analyzer (Seahorse Bioscience, North Billerica, MA). XFp Glycolysis Stress Test Kit (Seahorse Bioscience, Part# 103017-100) was used to detect three indexes including Glycolysis, Glycolytic Capacity and Glycolytic Reserve. Before the experiments, shNC cells or shNEAT1v2 cells were seeded at 5000 cells/well in 8 wells Seahorse cartridge which was hydrated before. After adhering, the medium in the wells was gently changed by supplementing XF Base Medium, which was added 2mM glutamine, as a starting point. The 8 wells Seahorse cartridge was incubated at 37°C in non-CO2 incubator for 1h. And then, the assay was performed through injection of 3 metabolic compounds: glucose, oligomycin and 2-DG. All the results were calculated on the average of the three measurements.

### Measurement of intracellular pH (pH_i_)

Measurements of pH_i_ were performed using the cell permeable pH-sensitive dye BCECF-AM (2’,7’-bis(2-carboxyethyl)-5(6)-carboxyfluorescein-acetoxymethylester; Calbiochem). Fluorescent probe BCECF (Aladdin, B115503) which is excited by visible light (488 nm) and diffuses out of cells very slowly was dissolved in DMSO for stock solutions (100mM), and the working concentration was 100nM. After adding BCECF to the culture medium and incubated in 37 °C for 20min, the flow cytometry analysis was performed to determine the intracellular pH of treated cells (BD Biosciences, Franklin Lakes, NJ, USA).

### Reagent treatment

Sirtinol (Sigma, S7942), a pan sirtuin inhibitor, was dissolved in ddH2O for stock solutions (2M). The cells were plated and treated with 10mM and 20mM sirtinol for 24h. Metformin (Abcam, #ab120847) was dissolved in ddH2O for stock solutions (1M) and used fresh or before 6 months. And subsequently diluted to different final concentrations: 0, 0.25, 0.5, 1, 1.5, 2, 4, 8, 10, 12, and 15mM with culture medium. C646 (Abcam, #ab146163) was dissolved in DMSO for stock solutions (20M) and diluted to different concentrations with culture medium.

### Statistical analysis

All assays were repeated at least three times and data are shown as means ±SEM. P values were determined by comparing the data from treated and control cells. Data were evaluated with two-tailed t-test. Differences were considered significant with a value of P<0.05.

## Acknowledgments

We are thankful to Pro. Naihan Xu and Pro. Weidong Xie for helpful discussion and critical reading of the manuscript. The authors thank the Graduate School at Shenzhen, Tsinghua University and funding supports from China government.

## Author Contribution

Yiwan Zhao and Ziqiang Wang designed and conducted the experiments, Yiwan Zhao analyzed the data, and wrote the manuscript. Yiwan Zhao, Yunhao Mao, Shikuan Zhang, Songmao Wang, Yuanchang Zhu, conducted the experiments. Bing Li conducted the bioinformation analysis. Weidong Xie and Naihan Xu revised the manuscript. Yaou Zhang designed the experiments, supervised the project, and wrote the manuscript. All authors read and approved the final manuscript.

## Appendix information

**Figure S1.**
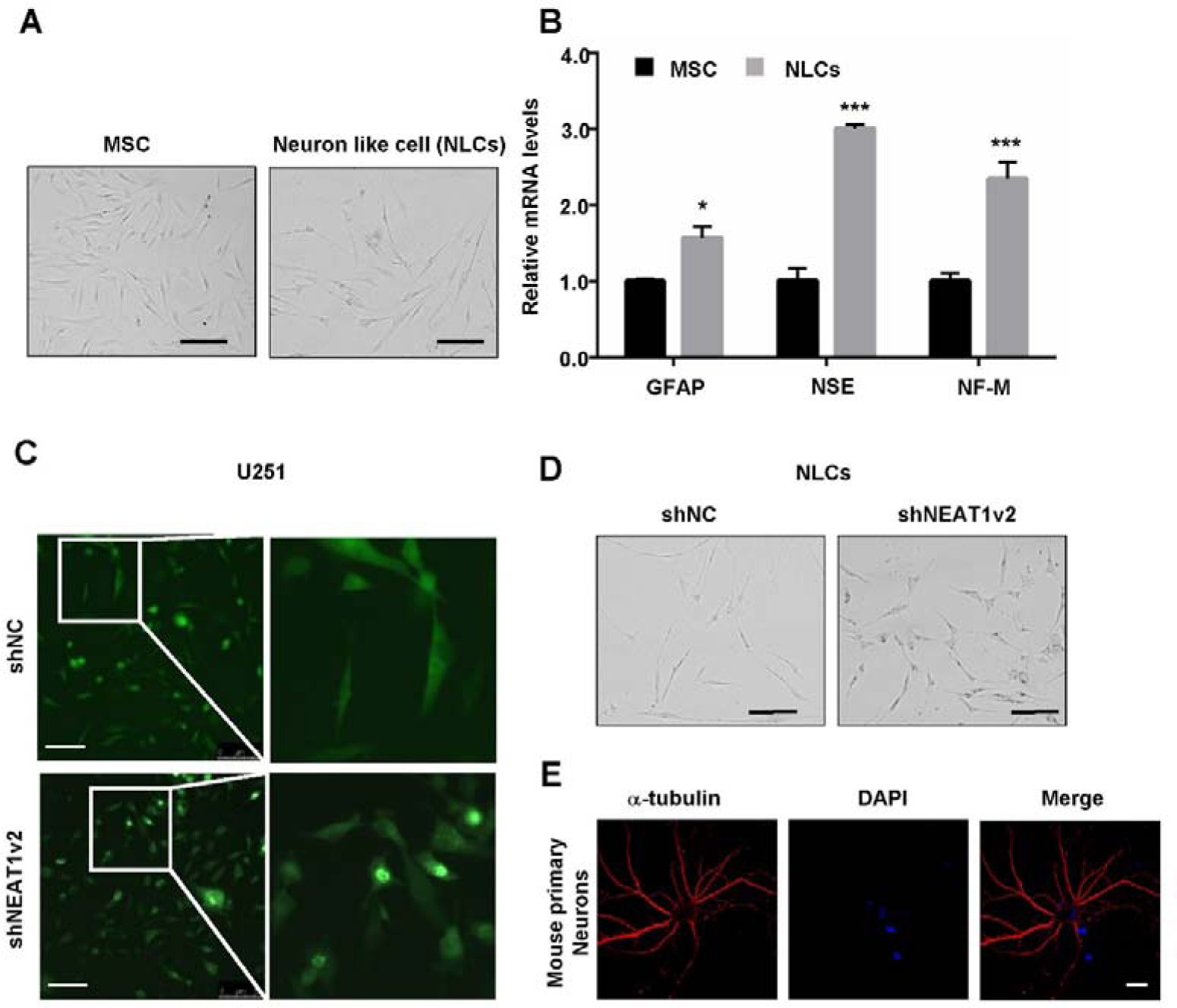
NEAT1 knock-down shorten the length of axon in Neuron like cells and synapse in U251. A. Morphologic change was observed after 4-6 day’s induction from MSCs to NLCs. Scale bars 100 μm. B. Quantitative PCR analysis of expression levels of several neuronal marker genes, GFAP, NF-M and NSE in MSCs and NLCs. C. The morphologic changes of stable cell line on U251 transfected with lentivirus based shRNA expressing green fluorescent protein (GFP). D. Shortened axonal length was observed after NLCs transiently infected with Lentivirus carrying shNEAT1v2 for 96h compared with shNC. Scale bars 100 μm. E. Immunofluorescence analysis of α-tubulin (red) in primary mouse neurons, DAPI (blue) was used to stain the nuclei. Scale bars, 20 μm.

**Figure S2.**
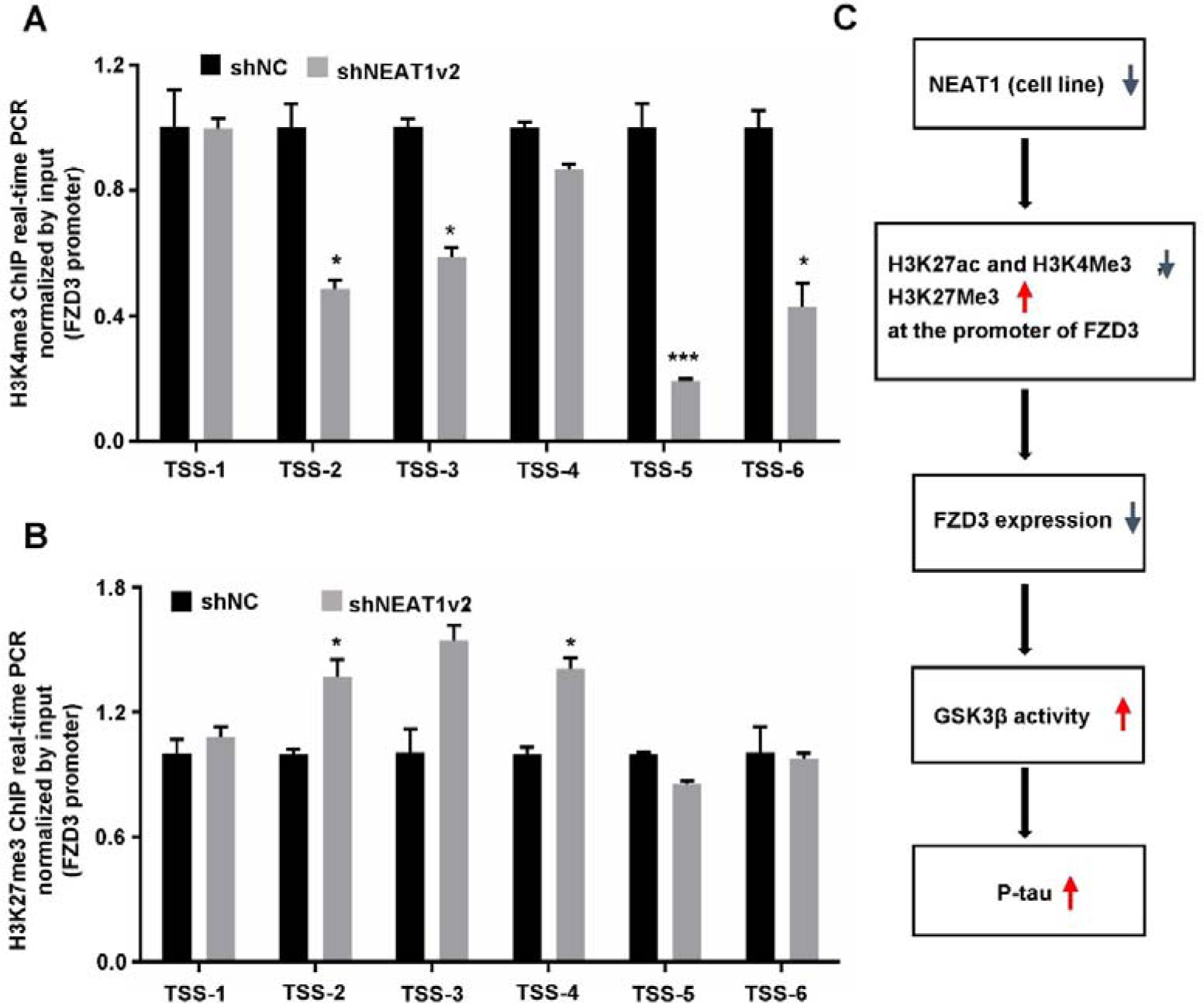
NEAT1 knock-down regulated FZD3/ GSK3β / p-tau signaling pathway via changes of histone modification. A. ChIP assays were performed with an anti-H3K4me3 antibody in stable NEAT1 deficient cell lines on U251, the fold enrichment of TSS1-6 fragments within FZD3 promoter by H3K4me3 relative to the input level was examined with real-time PCR. B. ChIP assays were performed with an anti-H3K27me3 antibody in stable NEAT1 deficient cell lines on U251, the fold enrichment of TSS1-6 fragments within FZD3 promoter by H3K27me3 relative to the input level was examined with real-time PCR. (mean ± s.d. of 3 independent experiments). \**P* < 0.05, Student 2-tailed *t* test. C. NEAT1 knock-down up-regulated phosphorylation of tau via inhibiting FZD3 expression and activated GSK3β signaling pathway.

**Figure S3.**
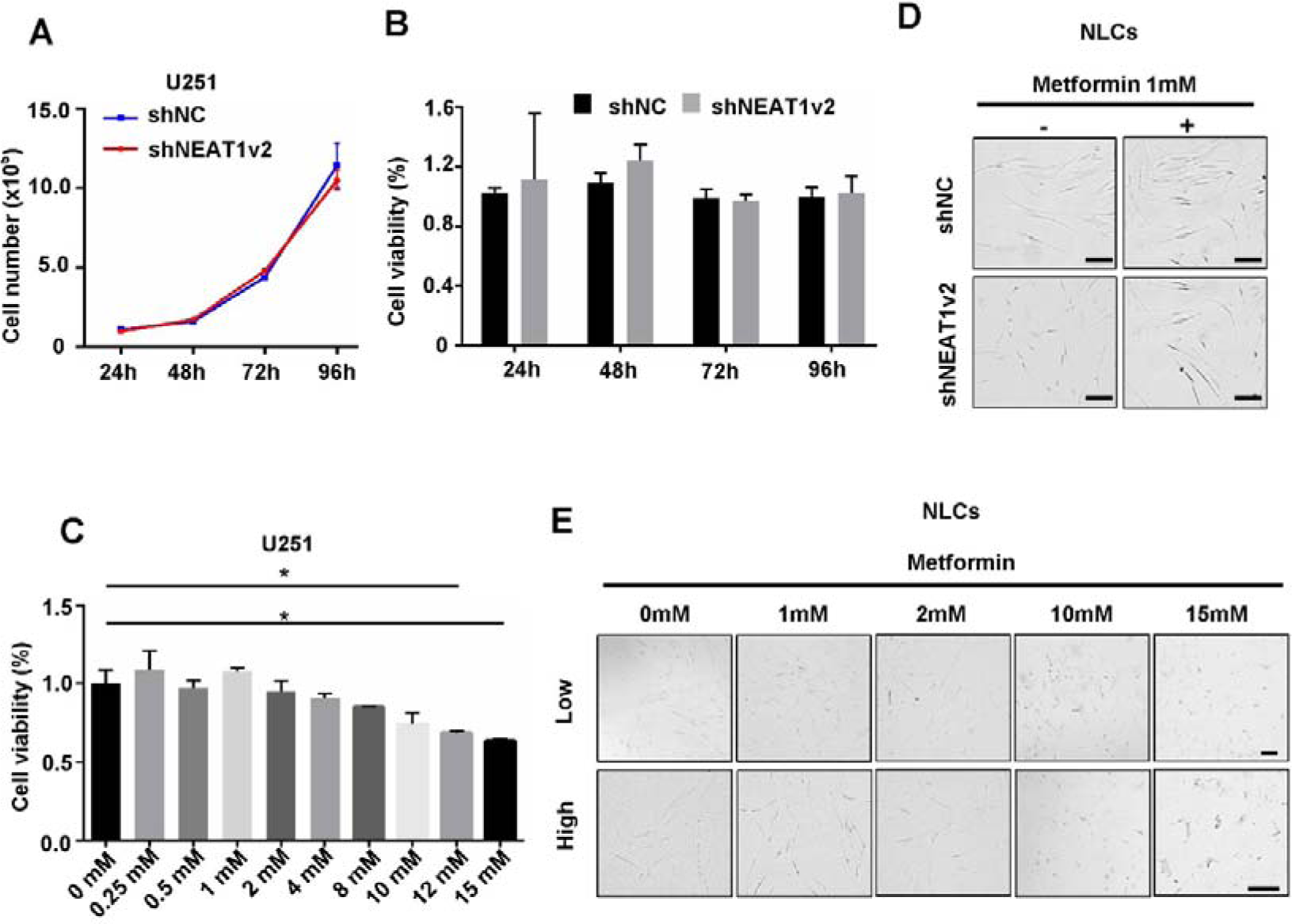
Several concentrations of metformin treatment influenced NEAT1 and FZD3 expression in U251 cells. A. The cell number of stable NEAT1 deficient cell line was analyzed by Hemocytometer after cultured 24h, 48h, 72h, and 96h in a 6-well plate. B. The cell proliferation was determined by Cell Counting Kit-8 at OD 450 nm in stable NEAT1 deficient cell line after cultured 24h, 48h, 72h, and 96h. C. The cytotoxicity of different metformin concentration (from 0 to 15mM) on U251 cells was determined by Cell Counting Kit-8 at OD 450 nm after 48h treatment. D. Rescued axonal length was observed on shNEAT1v2 and shNC transfected NLCs after treated with 1mM metformin for 48h under optical microscope. Scale bars 20 μm. E. The U251 cells were cultured with 1mM, 2mM, 10mM and 15mM metformin for 96h. Cell lesion could be observed in 10mM and 15mM metformin treated wells under optical microscope. Scale bars: 100 μm (mean ± s.d, \**P* < 0.05, Student 2-tailed *t* test).

**Table S1.**
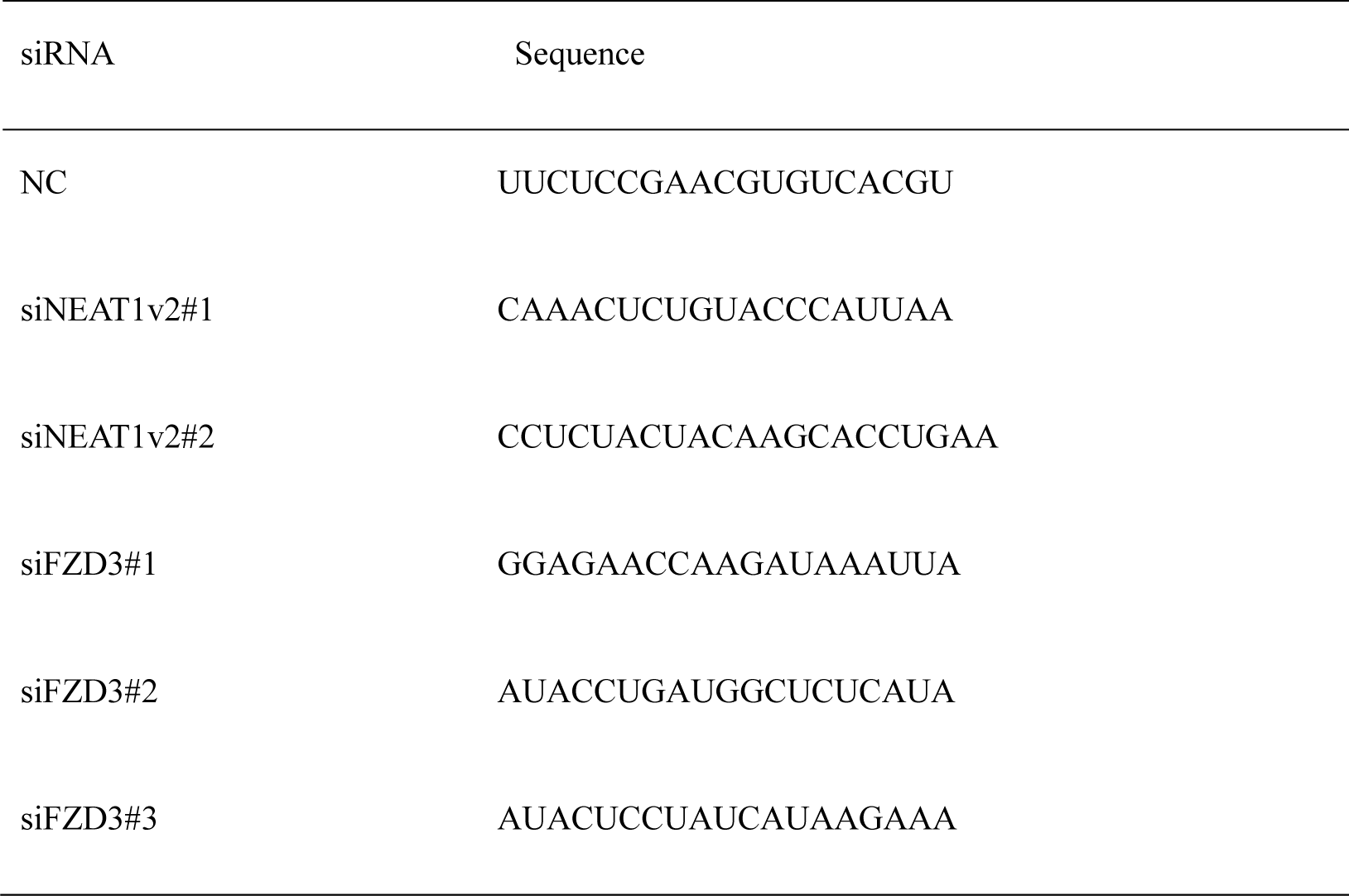
List of siRNAs sequence.

**Table S2.**
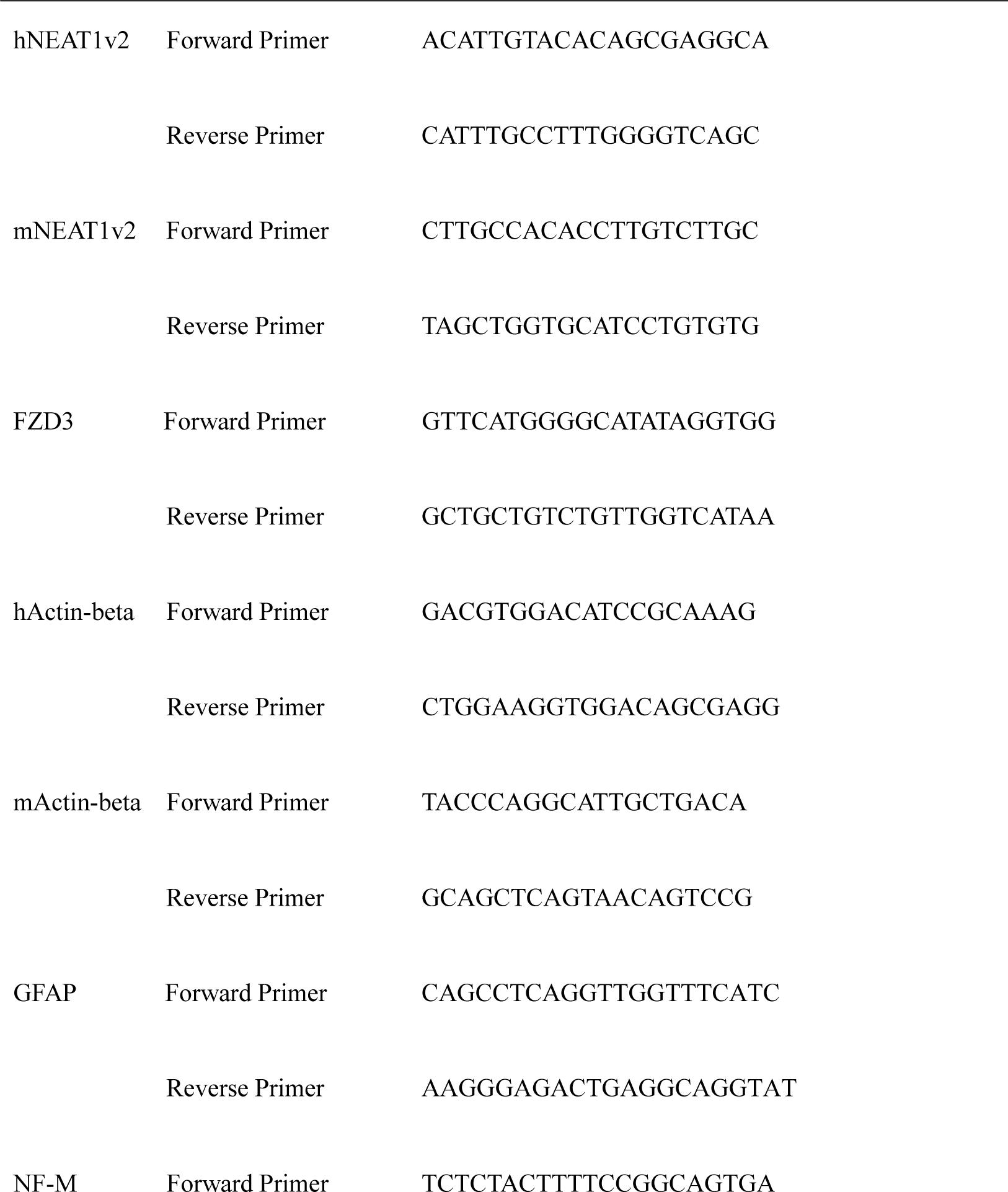
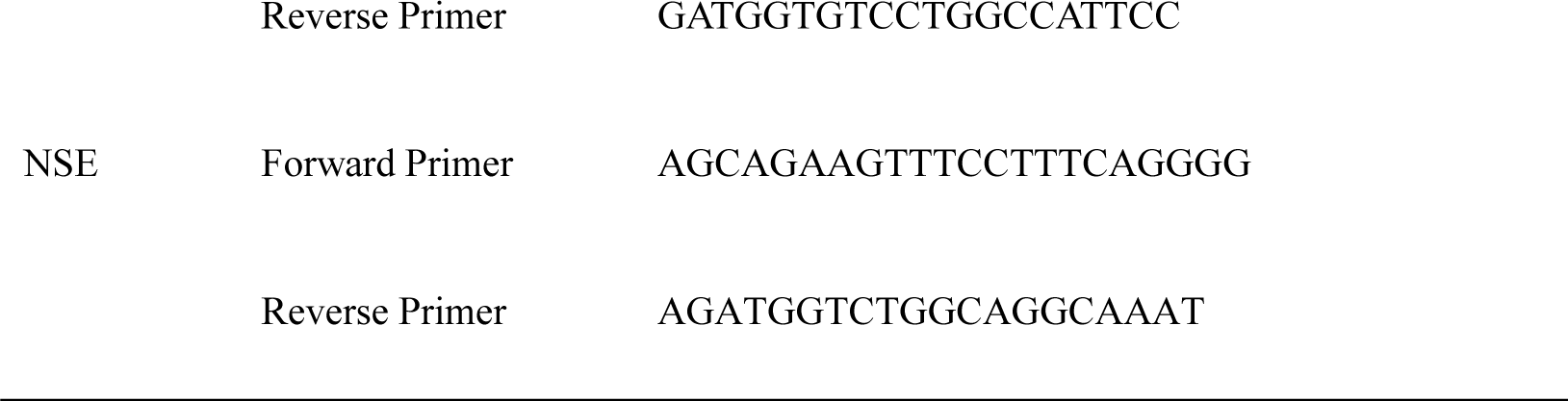
List of primers used for RNA analyses.

**Table S3.**
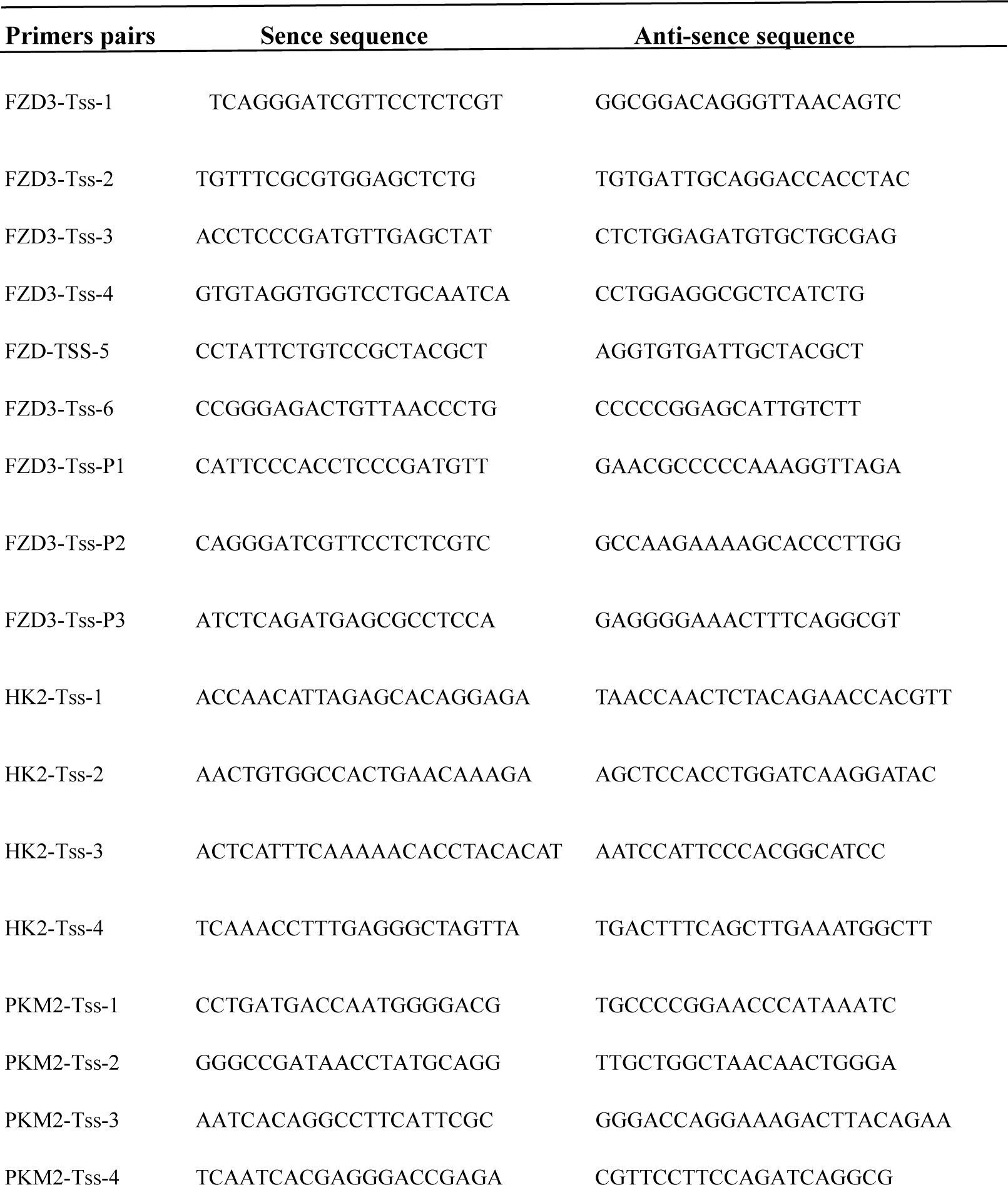
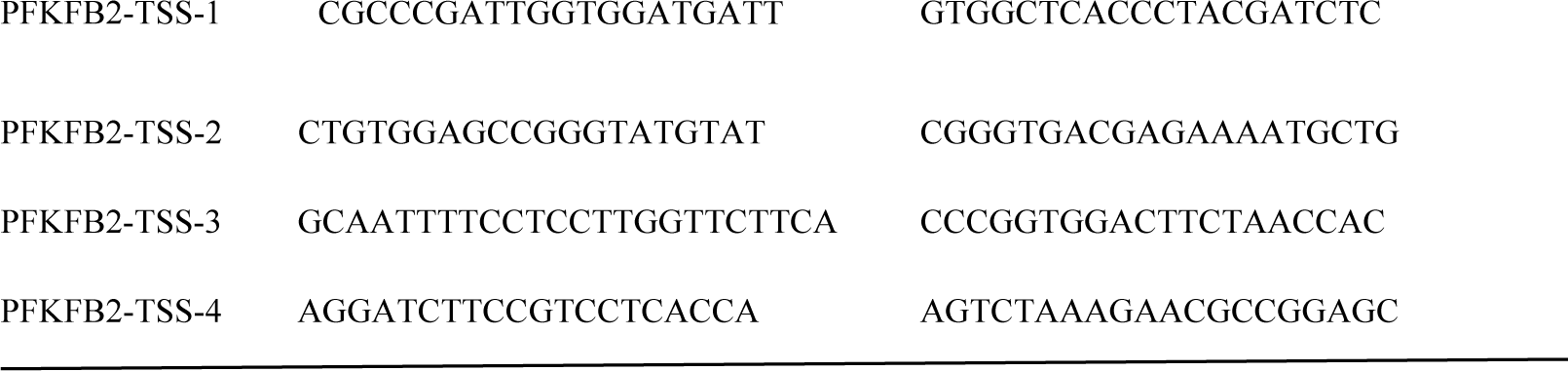
List of primers used for ChIP analyses.

